# Do maternal antibodies facilitate hemagglutinin imprinting to influenza A viruses encountered early in childhood?

**DOI:** 10.1101/110981

**Authors:** Katelyn M. Gostic, Monique Ambrose, Michael Worobey, James O. Lloyd-Smith

## Abstract

In a recent study, we showed evidence for childhood HA imprinting, a phenomenon in which children develop preferential, lifelong immune memory against zoonotic influenza viruses with hemagglutinin (HA) antigens in the same phylogenetic group as the first influenza virus encountered in childhood (Gostic et al. 2016). Although our original study showed strong, population-level HA imprinting effects, it did not resolve the underlying immunological mechanisms.

Similar immune imprinting phenomena, where individuals preferentially recall immune responses primed early in life, also influence seasonal influenza epidemiology via antigenic seniority (Lessler et al. 2012) and original antigenic sin (Francis 1960), yet the mechanisms underlying all these childhood immune imprinting phenomena remain poorly understood (Cobey & Hensley 2017). A recent letter from Dr. H. Lemke (Lemke 2017) suggested that these childhood imprinting effects might be mediated by the combined action of maternal antibodies (mAbs) and influenza antigen. In other words, that imprinting may require that children are exposed to influenza A virus in the first year of life, while maternal antibodies are still present.

## Introduction

As we responded elsewhere (Gostic et al. 2017) this hypothesis seems qualitatively inconsistent with epidemiological and virological data on influenza. However to be thorough, we performed model comparison to test formally whether epidemiological data better supported our originally published models (where primary exposure at any age from 0 to 12 years could induce HA imprinting) or two alternative models that include maternal effects. In **maternal effects model 1**, we assumed that imprinting could only occur in the first year of life. In **maternal effects model 2**, we assumed that primary influenza exposures throughout childhood could induce HA imprinting (as in the original study), but primary exposures in the presence of maternal antibodies might enhance imprinting protection. Here, we report the results of these quantitative analyses, and our methods, and show that the addition of maternal effects does not improve our existing models' predictions of birth year-specific pandemic influenza risk.

## Results and Discussion

After fitting all models to H5N1 and to H7N9 data, our original models performed better than either of the new maternal imprinting models (Tables 1 and 2). (Relevant data on case age distributions of emerging influenza viruses H5N1 and H7N9 are described in (Gostic et al. 2016).) Maternal effects model 1, which assumes HA imprinting can only occur in the presence of maternal antibodies (i.e. if primary exposure occurs in the first year of life) was definitively worse, with ΔAIC of 59 when fit to H5N1 data and ΔAIC of 80 when fit to H7N9 data. Thus, the original model in which HA imprinting could arise from primary influenza exposures throughout childhood (age 0-12) was much more consistent with observed epidemiological patterns than the alternate model in which imprinting could only arise while maternal antibodies are still present.

**Table 1:**
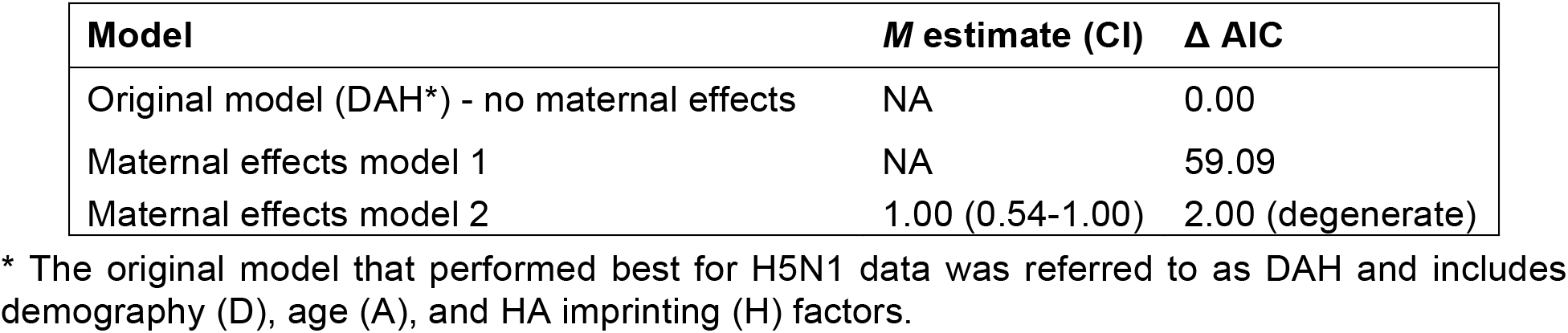
Model comparison results after fitting to H5N1 data.

**Table 2:**
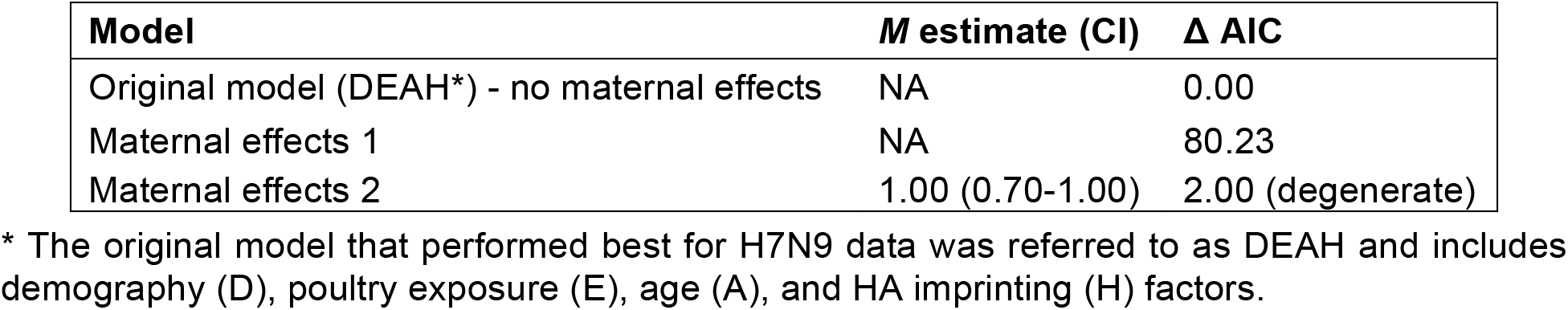
Model comparison results after fitting to H7N9 data.

In maternal effects model 2, where maternal effects might enhance imprinting, parameter *M* quantified the additional protection from maternally enhanced HA imprinting, with *M* = 0 indicating full protection and *M* = 1 indicating no signal of additional protection from imprinting in the first year of life, relative to imprinting later in childhood. Whether we fit the model to H5N1 or H7N9 data, the maximum likelihood estimate of *M* was 1, indicating no detectable effect of maternal enhancement. Because maternal effects are estimated to have no effect, model 2 is “degenerate.” In other words, it represents an algebraically identical, but more complicated version of the original model. The addition of parameter *M* simply multiplies the estimates from the original model by an extraneous factor of 1, and adds no new information. The original model and model 2 thus have identical likelihoods, but the original model requires one less parameter to produce the same estimates. Thus, maternal effects model 2 has ΔAIC of 2, reflecting the two-point penalty for each additional model parameter.

In summary, the original model (no maternal effects) is definitively the best model in this set. Upon fitting to both H5N1 and H7N9 data, the addition of maternal effects either decreases the model's ability to reproduce observed risk patterns (as in model 1), or complicates the original model without providing any additional information (as in model 2). Including maternal effects did not improve upon our original model's ability to estimate a particular birth cohort's epidemic or pandemic risk. The lack of statistical support for models in which mAbs are necessary to induce HA imprinting, or in which imprinting in the presence of mAbs enhances protection, suggests that HA imprinting and other childhood immune imprinting phenomena are driven by immune mechanisms unrelated to maternal antibody effects (Cobey & Hensley 2017).

## Methods

### I. Brief summary of previously published methods (Gostic et al. 2016)

Serological and epidemiological data show that the annual probability of influenza A virus (IAV) exposure is about 30% in naïve children, and that almost all children have had their primary exposure by age 12 (Sauerbrei et al. 2014; Sauerbrei et al. 2009; Neuzil et al. 2002; Bodewes et al. 2011). Thus, to estimate the fraction of each birth year with primary exposure (i.e. HA imprinting) to a group 1 or group 2 IAV, our original model took a weighted average of primary exposure probabilities across the first 13 years of life (from age 0-12). Here, the overall probability that someone born in year *i* was first exposed to IAV subtype *s* depended on:

- *ε_ij_*, the probability that an individual born in year *i* had their primary IAV exposure in year *j.*
- *f_s|j,c_*, the fraction of IAVs of subtype s that circulated in the year of primary exposure *j*, and country of residence, *c.*

To calculate *w_s,i|y,c_*, the fraction of birth year cohort *i* with primary exposure to subtype s, given data collected in year *y*, country *c*, we summed across all possible years of primary exposure, starting with the year of birth *(i)*, and continuing up to the first of two possible stopping points: the year in which the data were collected (*y*), or 12 years after birth *(i+12).*

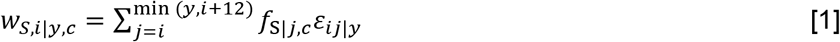

This is equation 6 in the methods section of the original publication (Gostic et al. 2016).

### II. Summary of new maternal effects models

In response to the letter from Dr. Lemke, we tested two new models in which maternal immunity could have different effects on HA imprinting:

**Model 1.** Assume that primary IAV exposure in the presence of maternal antibodies (during the first year of life) **is necessary to induce HA imprinting**.
**Model 2.** Assume that primary IAV exposure throughout childhood (age 0-12) can induce HA imprinting, but primary exposure in the presence of maternal antibodies (during the first year of life) **enhances HA imprinting protection**.

In both cases, we assumed that the presence of any maternal antibodies during the first year of life could induce HA imprinting, irrespective of which IAVs the mother had been exposed to, or at what points in her life those exposures took place. It is possible that the proposed role of mAbs in HA imprinting could be mediated by the abundance of mAbs present in the child, and by maternal exposure to an IAV of the same HA group encountered by the child. Thus, the full history of exposure to different IAV subtypes throughout the mother's lifetime might be important. To complicate matters even further, scientific arguments could be made to support a number of conflicting ideas about which maternal IAV exposures would have the strongest influence on her child's HA imprinting. Perhaps a mother's IAV exposures during childhood (Gostic et al. 2016; Cobey & Hensley 2017), throughout life (Lemke et al. 2004), or during pregnancy or breastfeeding (Okamoto et al. 1989; Lemke et al. 1994) would be most influential. Distinguishing between these hypotheses would require a full-scale research effort and is beyond the scope of the analyses presented here. Targeted experiments would provide the most power to distinguish among possible maternal imprinting mechanisms. Thus, in the absence of clearer mechanistic guidance from the literature, we made the simplest and most parsimonious assumption, in which any primary IAV exposure in the first year of life is sufficient to induce or enhance imprinting by the combined action of mAbs and IAV antigen.

### III. Estimating the probability of primary exposure in the first year of life vs. later in childhood

Models 1 and 2 require estimates of the fraction of each birth year whose primary influenza exposure occurred in the first year of life, as opposed to later in childhood. To estimate these quantities, we partitioned the above *w_s,i|y,c_* estimates as follows:

Define *x_s,i|y,c_* as the probability that a case born in year *i*, country *c*, and observed in year *y* had **primary exposure in the first year** of life to subtype *s*.

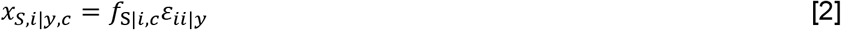

Define *Z_S,i|y,c_* as the probability that a case born in year *i*, country *c*, and observed in year *y* had **primary exposure in the 2^nd^**-**13^th^ year** of life to subtype *s*

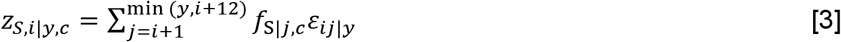

Note that equations 2 and 3 are modifications of equation 1, where the only difference is that we sum across different years of possible primary exposure *(j)*. Also note that *w_s,i|y,c_* = *x_s,i|y,c_* + *Z_S,i|y,c_*.

### IV. Model definition of birth year-specific risk of severe infection

Each model in our original analysis aimed to estimate the probability *p_yci_* that a given case observed in country *c* and year *y* occurred in an individual born in year *i.* The best-fit model defined these birth year-specific probabilities of case observation as:

Best model for H5N1 data (DAH):

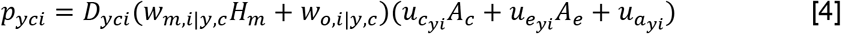

Best model for H7N9 data (DEAH):

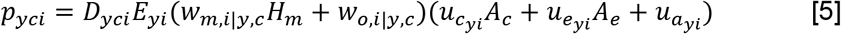

Here, *D_yci_* represents the fraction of the population of country *c* in year *y* that was born in year *i. E_yi_* represents the age-specific risk of contact with the poultry reservoir. Parameter *H_m_* describes the relative risk of severe infection in individuals with protective HA imprinting, and modulates *w_m,i|y,c_*, the fraction of birth cohort *i* with protective imprinting (primary exposures <u>m</u>atched to the challenge strain). *w_o,i|y,c_* is the complement of *w_m,i|y,c_*, and gives the fraction of birth cohort *i* with non-protective imprinting (<u>o</u>ther primary exposures). Parameters *A_c_* and *A_e_* describe relative risk of severe infection in individuals belonging to high-risk age groups (children age 0-4, and elderly adults age 65+, respectively). All *u_x_* are binary indicator variables representing whether each case in the dataset belongs to the childhood high-risk age group, elderly high-risk age group or baseline adult age group. Variables and parameters are described in greater detail in the supplementary text of the original article (see section S2), and in table S1 (Gostic et al. 2016).

For maternal effects model 2 we defined a new parameter, *M.* Similar to *H_m_* in the original models, *M* represents the relative risk of severe infection in individuals protected by maternally enhanced imprinting, relative to all others. *M* takes values from 0-1 with 0 representing complete protection and 1 representing no detectable protective effect beyond that represented by *H_m_.*

Below, in equations 6 and 7, subscripts for fractions *x* and *z* (as defined in equations 2–3) follow the same conventions as overall imprinting weights (*w*), where subscript *m* indicates group <u>m</u>atched (i.e. protective) primary exposure, and subscript *o* indicates <u>o</u>ther (non-protective) primary exposure:

Maternal effects model 1 (Primary exposure in the presence of maternal antibodies **required** for HA imprinting)

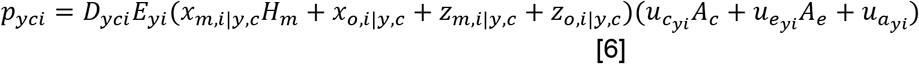
Maternal effects model 2 (Primary exposure in the presence of maternal antibodies **enhances** HA imprinting protection)

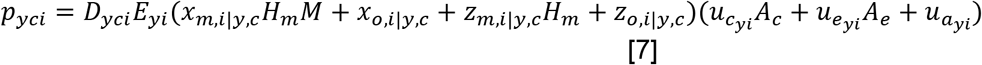

Note that each of these models is an extension of the DAH/DEAH framework presented above.

We fit each of these models and the original models to H5N1 and H7N9 data to estimate maximum likelihood values of all relevant parameters, and to find profile confidence intervals, as described in (Gostic et al. 2016). We then performed model comparison (using AIC) to determine which model best described the data. Note that in the original study, factor *E_yi_* was included in the best-fit model for H7N9 data but not for H5N1. Thus, we only included this factor when fitting to H7N9 data in this analysis, and excluded it when fitting to H5N1 data

